# Decision bias and sampling asymmetry in reward-guided learning

**DOI:** 10.1101/2023.09.10.557023

**Authors:** Yinan Cao, Konstantinos Tsetsos

## Abstract

Human economic decisions are highly sensitive to contexts. Deciding between two competing alternatives can be notoriously biased by their overall value (‘magnitude effect’) or by a third decoy option (‘distractor effect’). Some prominent explanations appeal to diminishing value sensitivity and divisive normalization in value representations, i.e., representational bias, that feed into the choice stage. However, these explanations have recently come under scrutiny due to empirical inconsistencies and mounting alternative theories. Here, we posit that context-sensitive choices may not stem from representational biases but rather emerge as by-products of asymmetric sampling during value learning. In a reward-guided choice task, participants aimed to maximize cumulative rewards through trial and error. The task introduced alternating blocks with either a colored distractor or a neutral ‘notional’ distractor. We observed decreased choice accuracy when higher-value distractors were present, a pattern that persisted even in the notional distractor blocks. Using computational modeling, we show that this phenomenon falls out naturally from a simple learning rule without relying on any additional mechanism such as divisive normalization or nonlinear utility. Furthermore, we found that, contrary to divisive normalization, choice accuracy was not influenced by distractor value but strongly depended on the magnitude of the targets’ values per se. This ‘magnitude sensitivity’ was also found in the ‘notional distractor’ conditions and could lawfully be reproduced by the learning model. Importantly, when counterfactual feedback eliminated sampling asymmetry, the observed decision bias vanished. Our results suggest that the genesis of context-sensitive choices may lie in the learning dynamics themselves, specifically sampling asymmetry, rather than in pre-decisional representational biases. This finding reframes the discourse on irrational decision-making, attributing it to acquired biases during the learning process, not necessarily computational intricacies at the choice stage.

## Introduction

Faced with a complex and uncertain world, how do we decide and act? This inquiry fuels a wide array of disciplines, ranging from neuroscience and economics to philosophy. Our decision-making process is shaped by the way decision-relevant information is sampled and encoded in neural systems. Imagine visiting a new city. Initially, we venture into various cafes and restaurants to explore the culinary landscape. However, upon discovering a particularly enjoyable restaurant, there is a tendency to exclusively frequent that establishment or a small set of safe options. Consequently, we forgo a holistic grasp of the city’s varied culinary offerings, a situation that exemplifies the classic explore-exploit trade-off^1–5^. Using an analogy in machine learning, once the optimal action is pinpointed, an artificial agent halts the process of sampling alternatives since acquiring precise values for those alternatives becomes less significant^6–8^. This exemplifies under-sampling as a pervasive ‘bias’ in biological and artificial systems, where certain choices are disproportionately favoured due to limited exposure during learning.

Akin to leveraging sensory illusions to understand the workings of the visual system, probing into the computational and neural basis of decision biases can shed light on the generic computations that underlie decision-making^11,12^. Humans and other animals appear to make biased reward-guided choices that violate axiomatic rationality^13,14^, e.g., deciding between two competing target economic alternatives can be swayed by a third distractor prospect^9,15–19^ (Fig. 1a, b). To elucidate this value-based ‘distractor effect’, some prominent explanations appeal to ‘divisive normalization’—a canonical neuronal computation originally uncovered within sensory neuroscience^20,21^—in value representations that later feed into choice stage^9^ (Fig. 1c). One challenge is that the processes governing decisions about sensory inputs and economic prospects can be quite different. In sensory information processing, the parameters shaping a percept typically adhere to certain transduction functions based on available sensory inputs. For economic decisions, by contrast, the pertinent parameters (e.g., the value of a consumer product or the pleasantness of a dining experience) need to be *learnt* through sampling of information, entailing their encoding in a utility function^10,22^ or retrieval from a model of the world^7,23^. A recent large-scale replication failure of the value-based distractor effect has spurred a heated debate over its empirical robustness^24–26^, casting doubt on the theory of divisive normalization at the level of value representation as the primary explanation^27–29^.

**Figure 1.**
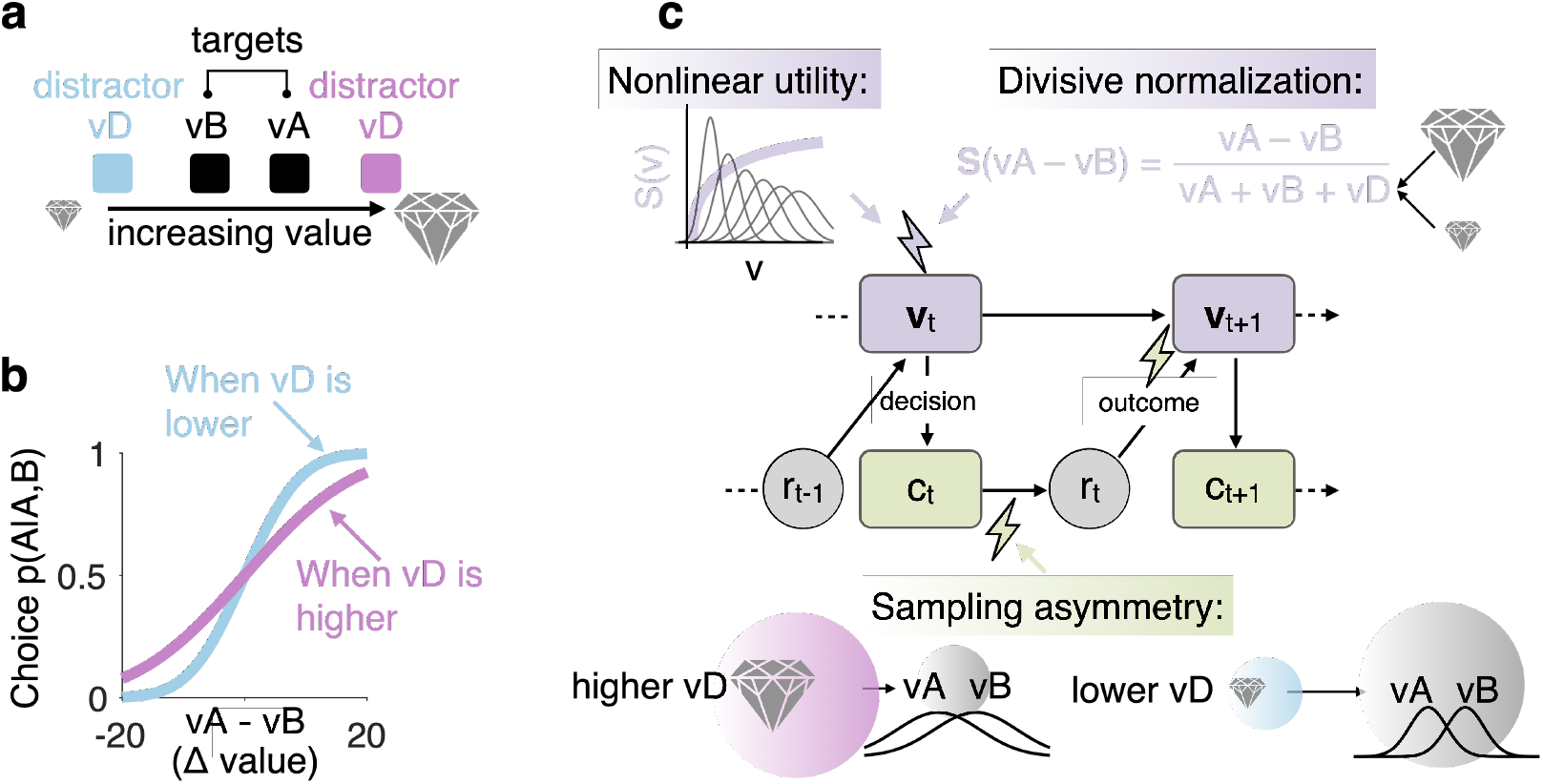
Value-based context effect and computational theories. (a) Value-based choice scenario. (b) Value-based context effect: Relative accuracy (conditional probability) of choosing the higher-value target is reduced by a higher-value distractor (shallower psychometric function). (c) Rival computational accounts for value-based context effect. Value-based divisive normalization^9^: a lower ‘subjective’ (S) value difference (the decision variable) between the two targets in the presence of a higher-value distractor due to a larger denominator. Divisive normalization operates at the stage of latent value ‘v’ before the normalized value representations are fed into the choice stage ‘c’. Moreover, value representation could be distorted by diminishing marginal utility, as exemplified by the power-law mapping between objective and subjective values^10^. Sampling-based learning asymmetry: A higher-value distractor (D) leads to increased exploitation of this option, or ‘over-sampling,’ thus causing the target options A and B to be relatively under-sampled. The size of the disk represents the frequency of sampling, indicative of participants’ propensity for exploitation or ‘greediness’ in choosing the option to learn its value and maximize overall payoff. This results in compromised value updates (wider/noisier econometric tuning curves) for A and B in comparison to D. The effects of sampling-based learning asymmetry iteratively influence subsequent trials and propagate forward into future decisions.

Concurrently, a lively debate has erupted over the potential inaccuracies in value coding across different contexts. Utility functions, for instance, frequently display a concave shape that aligns with the concept of diminishing marginal utility, a key tenet of expected utility and prospect theory^10,30^. These functions introduce a power-law relationship to convert objective measures into subjective utilities^22^. When examining magnitude or numerical values, the structure of econometric functions becomes more intelligible if one assumes a log-normal, or otherwise compressive, encoding^31^. Such an assumption broadens the tuning curves for larger values, leading to less precise estimates. This compressive feature of the utility function sheds light on why overall magnitude—which essentially equates to the mean value of available choices—influences decision-making performance. This specific influence is known as ‘magnitude sensitivity’^32^. Notably, in high-magnitude scenarios, decision-makers tend to respond both more quickly and more erratically^33^, although cases of the opposite behavior have also been reported^34^.

Of significant note, the way reward information is sampled plays a direct role in shaping value representations, as illustrated in the ‘restaurant sampling’ example. Prior investigations into the distractor biases in reward-driven decision-making have often neglected to explicitly consider the effects of learning and information sampling. This oversight bears heightened relevance, particularly considering that a substantial number of human studies, and quite possibly the majority if not all studies involving animals, necessitate a training phase to ensure the decision-makers grasp the concept of value (usually mapped to abstract pictures or visual features^19,35^). Now consider a scenario where two target prospects, A and B, are paired with higher-or lower-value distractor prospects (as depicted in Fig. 1a) across two distinct experimental blocks. As decision-makers learn these prospects’ values through trial and error, a biased preference for the higher-value distractor could lead to the under-sampling of target options (Fig. 1c). This, in turn, may undermine the value representation for the two target prospects, leading to heightened uncertainty about their worth (wider econometric tuning curves; Fig. 1c), particularly when contrasted with the block where the distractor holds a lower value. This example highlights the idea that the distractor effect can occur as epiphenomena of sampling asymmetries during reward learning, a novel perspective that has not been previously explored.

Reward sampling asymmetries result in changes in the shape of the econometric tuning curve (as shown in Fig. 1c), which can be mistaken for representational value distortion, but these two processes are dissimilar in nature. The key difference between the effects of reward sampling asymmetry and that of value representational distortion—whether arising from a nonlinear marginal utility function or divisive normalization—resides in their respective loci of influence. Representational distortions take effect at the valuation stage itself, directly shaping perceived value. By contrast, sampling asymmetry affects the pathway from choice to outcome at each iterative update step and subsequently reverberates into future trials, thereby displaying a recursive and cascading nature. As such, these two mechanisms fulfill distinctly different computational roles and functions. However, it remains an open question whether contextual influences on value-based decision, such as magnitude sensitivity^32,34^ or distractor effect^9^, originate solely from representational biases at the pre-choice stage or may also arise from sampling asymmetry during value construction in the learning process.

Here, we developed an experimental framework in which the effects of sampling asymmetry and representational distortion on decision bias can be compared in the context of the same reward-guided choice task (Fig. 2a). In a baseline condition, participants learn to maximize their payoffs by repeatedly selecting a high-value target amidst a distractor that is present but un-selectable. We then crafted conditions where divisively normalizing option values using a larger distractor value is not feasible. We achieved this by making the distractor ‘neutral’ and non-informative regarding value, rendering it merely ‘notional’. This setup attributes any effect of a notional distractor exclusively to sampling asymmetry. Additionally, we neutralized sampling asymmetry by providing counterfactual feedback on both selected and unchosen options, thereby attributing any observed magnitude sensitivity in behavior solely to representational bias. We found both distractor effects and magnitude sensitivity in the baseline condition: the accuracy in selecting a higher-value target diminished when faced with a higher-value distractor but improved when the target options had higher magnitudes. Importantly, these phenomena were also robustly observed in the ‘notional distractor’ condition. Through computational modeling, we demonstrate that these outcomes are naturally predicted by a simplistic learning model without the need for additional mechanisms like divisive normalization or nonlinear utility. Further, counterfactual feedback on unchosen options nullified the effect of sampling asymmetry, thereby eradicating the magnitude effect in choice. These findings indicate that context-sensitive choices, often considered a hallmark of irrational decision-making, may be attributed to biases developed over the course of learning rather than to computational processes occurring during pre-decisional deliberation.

**Figure 2.**
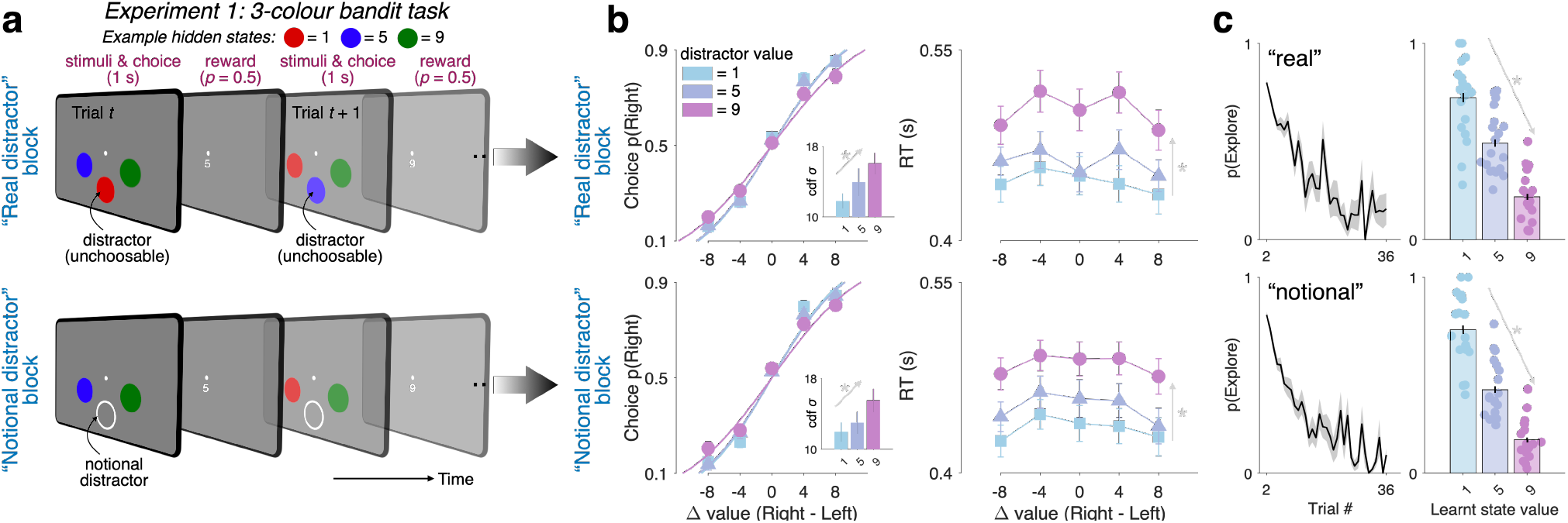
Task design and behavioral results from Experiment 1. (a) Each trial presented three iso-eccentric stimuli, uniquely color-coded as red, green, or blue. Flanking a central fixation point, two target stimuli were available for selection via button press, while a third ‘distractor’ stimulus, positioned below the fixation, was not selectable (no button offered). Colors were associated with fixed reward magnitudes (with a reward probability of 0.5), drawn *with repetition* from a set of values (1, 5, or 9 cents). Different colors could have the same reward magnitude (see Methods). The mapping of color to reward remained constant within each block but was fully counterbalanced across blocks (3 x 3 x 3 = 27 combinations). Participants were tasked with learning the value of each color through trial and error, aiming to maximize cumulative rewards. Crucially, alternating blocks featured either a colored distractor (‘real’) or a non-informative, white circle serving as a ‘notional distractor.’ (b) Probability of selecting the right-side option and the associated reaction time (RT), as functions of the value difference between the right and left options (value of right minus value of left) as well as distractor type. The standard deviation of a Gaussian cumulative density function (CDF sigma) was fit to the probabilities of choosing the right-side option (p(Right)). Differently colored curves and markers denote conditions where the distractor’s reward value was set at 1, 5, or 9 cents, with conditions aggregated across various learning blocks. (c) Probability of opting for an unknown, previously unchosen option, plotted either as a function of trial number (left panels) or as a function of the known option’s value that had been chosen previously (right panels). Upper panels present data from ‘real distractor’ contexts, while lower panels showcase data from ‘notional distractor’ contexts. Error bars = ± s.e.m. across participants (*N* = 20). Asterisks: significant effects (*p* < .05; 2-sided 1-sample *t-*tests against 0) following Holm’s sequential Bonferroni correction for multiple comparisons.

## Results

In a sequence of experiments using a modified multi-armed bandit paradigm, we investigated the influence of an ‘unavailable’ distractor stimulus on decision-making processes. In all experiments, participants were instructed to select between two visual stimuli positioned to the left and right of a central fixation point (Fig. 2a). Crucially, an additional stimulus, located at the bottom of the screen and explicitly designated as ‘unavailable for choice,’ served as a distractor. The presence of this distractor permitted us to systematically dissociate overt choice sampling from latent valuation, thereby clarifying its role in modulating decisions between available targets. To further elucidate the influence of reward learning dynamics on decision quality, the reward probability for chosen stimuli was fixed at 0.5, precluding one-shot learning.

### Pseudo-distractor effect occurs in ‘notional distractor’ contexts

In the first experiment, participants were trained to learn hidden rewards of three distinct colors— red, green, and blue (Fig. 2a). Each color was associated with a reward magnitude (reward p = 0.5), selected (with repetition) from a set of increasing values (1, 5, and 9 cents), and the color-reward mapping was fixed within a block but was fully counterbalanced across blocks. We constructed psychometric curves to quantify the accuracy of selecting the higher-value option as a function of the value difference between the two available targets. Participants showed an increased aptitude for selecting the higher-value option when the value difference between the two targets was pronounced (Fig. 2b). Interestingly, the slope of the psychometric curve was significantly modulated by the value of the distractor. As the distractor’s value grew from 1 to 5, and then to 9 points, the slope of the psychometric curve became progressively flatter (Fig. 2b). This flattening appears to suggest a reduced sensitivity to the value difference between the two targets when distractor value increased. A Gaussian cumulative density function (CDF) fit to the choice accuracy data confirmed that the standard deviation of the function expanded significantly (*p* < .05) with increasing distractor value (Fig. 2b; upper left panel). Additionally, reaction times (RT) correspondingly prolonged as a function of the distractor’s value (Fig. 2b; upper right panel).

Intriguingly, this ‘distractor effect’ persisted even in the ‘notional distractor’ blocks. In these blocks, the distractor’s color was replaced by a non-informative white circle, effectively serving as a ‘placeholder.’ Despite the absence of an informative distractor, the slope of the psychometric curve and RTs exhibited similar trends to those observed in the ‘real distractor’ blocks (Fig. 2b; bottom panels). This finding suggests that the distractor’s influence may not solely be attributed to its value but likely covaries with other, as-yet-unidentified modulators of choice accuracy.

### Pseudo-distractor effect emerges from asymmetric sampling

Next, we aimed to identify the latent variables that were influencing choice accuracy, variables that seemed to co-vary intriguingly with the distractor value. To delve into this, we examined participants’ inclination toward exploration. We quantified this exploration tendency by charting the probability of opting for an unexplored option against the value of an already selected but unchosen alternative. This measure of exploratory behavior serves as a proxy for the adaptive nature of decision-making strategies as learning progresses (Fig. 2c).

Our analysis yielded two notable trends. Firstly, as expected, participants’ willingness to opt for an unfamiliar option decreased as they accrued experience over a series of trials (Fig. 2c; left panel). Secondly, and of particular significance, the propensity for exploration was inversely correlated with the value of a known, yet unchosen, option (Fig. 2c; right panel). This suggests that participants curtailed their exploration of unknown options when they had already identified a high-value alternative in the past. Intriguingly, this pattern of exploration remained consistent across both the ‘real distractor’ and ‘notional distractor’ conditions, with no difference observed in either the magnitude or trend of exploratory behavior (Fig. 2c, upper vs. lower panels). This consistent behavior across conditions hints at the possibility that the observed decline in decision quality—particularly when selecting between two lower-value options (Fig. 2b)—is attributable to an asymmetrical learning process. Specifically, a high-value option, when overexploited, seems to impede the accurate learning of other options with lower intrinsic values.

These findings challenge the assumption that the observed distractor effect on choice accuracy is solely attributable to divisive normalization processes at the decision-making stage. This assumption would have predicted a significant disparity between the real and notional distractor conditions, which our data do not support. To substantiate our alternative explanation—that the effect stems from asymmetries in learning—we employed computational modeling using the delta rule, a classic model of reinforcement learning. This parsimonious model incorporates just two key parameters: a learning rate, which modulates the influence of reward prediction errors on updating latent value estimates for the available options, and a softmax temperature, which controls the degree of stochasticity in action selection relative to the value differences between the two target options—essentially accounting for action noise. Notably, the model does not encompass any mechanisms akin to divisive normalization, making it an ideal candidate for testing our hypothesis.

Upon fitting this model to human trial-by-trial choice data, we found that it well captured the learning dynamics, despite its minimal parameterization (Fig. 3a). Most compellingly, the model succeeded in replicating the distractor effects observed in both real and notional distractor conditions (Fig. 3b). This lends credence to our argument that the distractor effect is not solely the product of divisive normalization but may rather originate from learning asymmetries.

**Figure 3.**
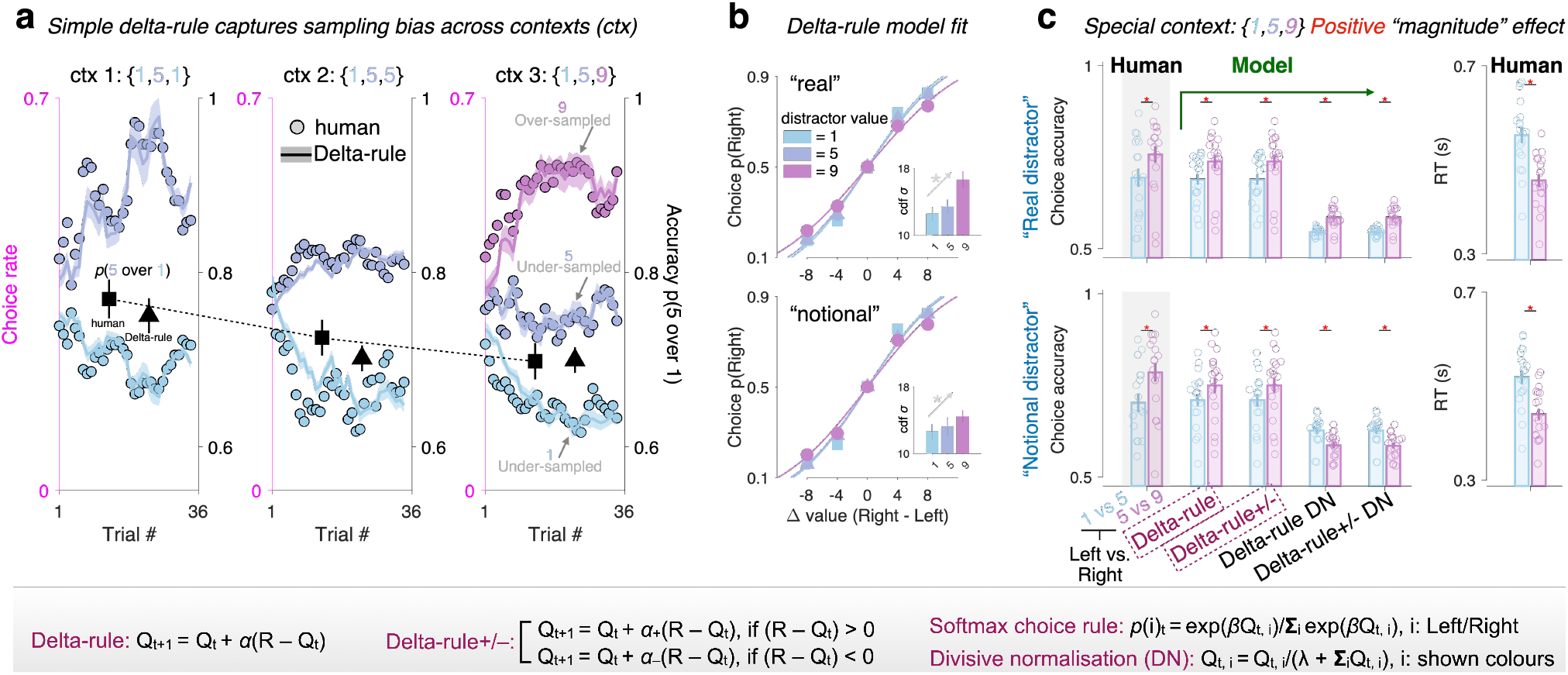
Simulated behavior by reinforcement learning model. (a) Illustration of learning curves, expressed through choice rate (averaged in a sliding window of 2 trials) as a function of option value and time (trial #). Three different contexts (ctx), or independent learning blocks, are shown for illustration. Across these contexts, two colors were assigned rewards of 1 and 5, while the third color received a reward of 1, 5, or 9. This design allows examination of how reward distribution across options affects learning and decision-making. The right axis (black) shows the mean accuracy for correctly choosing a higher-valued option (reward = 5) over a lower-valued one (reward = 1). Square: human; triangle: model fit. (b) Choice probability for the right-side option (p(Right)) as predicted by the learning model fit to human data. (c) Choice accuracy and RT for two conditions with the same value difference between targets (1 vs. 5 and 5 vs. 9) but different mean values. The equations at the bottom outline the learning model architecture. Q is the latent value estimate updated by a reward prediction error, R - Q, at each step t during learning. R is the outcome associated with a certain choice, α is the learning rate (ranging from 0 to 1) scaling the prediction error before updating Q. The parameter β in the softmax choice rule indicates the inverse temperature, with lower β implying higher action stochasticity. In divisive normalization, Q values are normalized by the total sum of displayed stimulus values. In the notional distractor condition, the denominator is the sum of the values of the two targets.

At first glance, it may seem counterintuitive that a distractor effect could manifest in the notional distractor condition. How could a non-informative placeholder wield any influence over the quality of decisions between two available targets? To elucidate this, we turn to Fig. 3a, where we illustrate how learning asymmetry engenders a pseudo-distractor effect across various learning contexts. We plotted the choice rates for different options as functions of their hidden values across three distinct learning blocks or contexts (‘ctx’). In each context, the three colors were assigned varying combinations of hidden values: 1, 5, and 1 point in the first; 1, 5, and 5 points in the second; and 1, 5, and 9 points in the third (Fig. 3a). Notably, in the third context where a high-value option (reward = 9) exists, participants’ propensity to repeatedly select this optimal choice markedly increased over the course of learning. This pattern resulted in the other two options being disproportionately under-sampled. Consequently, the choice accuracy for an option valued at 5 points over one valued at 1 point diminished as the learning environment shifted from ctx 1 to ctx 3, as illustrated in Fig. 3a (markers— black squares representing human behavior and black triangles indicating model predictions). This observed decline in choice accuracy underlines the asymmetric sampling induced by the presence of a high-value option, offering a cogent explanation for the pseudo-distractor effect seen in both real and notional distractor conditions.

### A positive magnitude effect naturally falls out of sampling asymmetry

Thus far, our analysis has demonstrated that a pseudo-distractor effect manifests across various learning contexts. This effect appears to arise due to an imbalanced exploitation of high-value options and the consequent under-sampling of lower-value alternatives. However, this does not entirely preclude the operation of divisive normalization within specific, localized context. To address this, we turned our attention to a unique context in which the three colors were associated with three distinct reward magnitudes: 1, 5, and 9 points (Fig. 3c). We scrutinized choice behavior under two conditions: when the value difference between the target options was held constant at 4, while only the value sum (magnitude) of the targets varied. According to the theory of divisive normalization, an increase in the magnitude of the target options—due to a larger denominator—should result in reduced decision accuracy. Intriguingly, observed human behavior contravened this prediction, aligning more closely with what a simple learning model would anticipate (Fig. 3c). This pattern remained consistent even when we introduced variant learning rates for positive versus negative reward prediction errors^36–38^, an adjustment commonly employed in previous research^39^. Importantly, a model that explicitly integrates a divisive normalization step^9^ at the choice stage produced inaccurate predictions, particularly in the notional distractor context (Fig. 3c, bottom panel). In this case, the normalization equation’s denominator solely comprises the values of the two target options, which led the model to forecast significantly lower overall decision accuracy than what was observed in human behavior. Additionally, we noted that RTs were faster when the value magnitudes of the targets increased. Collectively, these results cast doubt on the notion that divisive normalization serves as a tenable explanation for the behaviors observed in our study.

The results of Experiment 1 suggested that the pseudo-distractor effect is consistent across both notional and real distractor conditions, implicating sampling asymmetry as a key contributor given its generalizability across these distinct contexts (‘real’ and ‘notional’ distractors). However, one could argue that the colorless notional distractor might implicitly carry the value of the unshown color— especially considering that the 3 displayed colors were different from each other on each trial. To address this, we designed Experiment 2 with an enhanced paradigm. In this refined design, participants were tasked with learning the values of four different colors (Fig. 4a; values ranging from 10 to 99). This allowed for the pairing of the target options with one of two possible distractor colors, enabling us to keep the target options constant while manipulating only the distractor value. This setup served as a stronger test to disentangle the influences of learning asymmetry from those of divisive normalization mechanisms. Our data indicated an increase in choice accuracy with elevated target magnitude, replicating the magnitude sensitivity observed in Experiment 1. Strikingly, we found no discernible impact of distractor value on decision accuracy, regardless of whether the distractor was real or notional (Fig. 4b, c).

**Figure 4.**
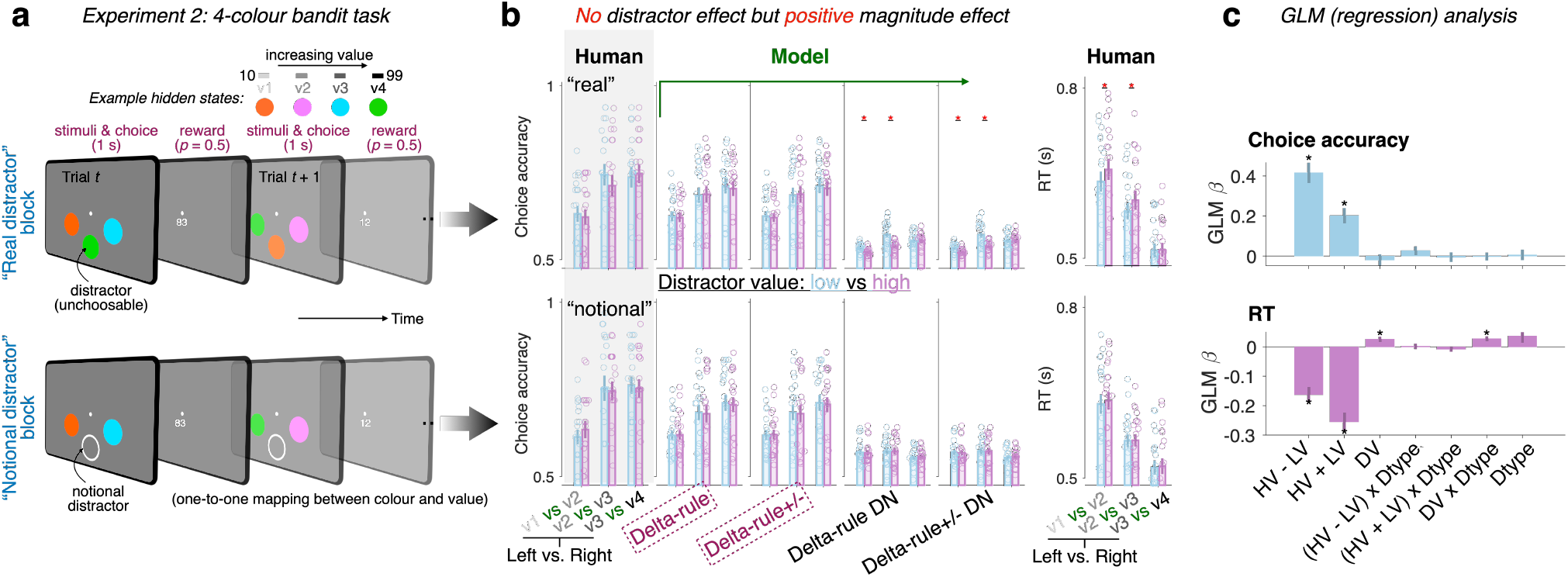
Task and behavioral data in Experiment 2 and simulated behavior by learning model. (a) Four colors were mapped to four non-overlapping, increasing reward ranges (Methods). As in Experiment 1, participants were required to choose between left and right targets on each trial, with the bottom distractor being unavailable for choice. Each trial featured three different colors, with no repeated colors among the three options. Across trials, stimulus locations and color combinations were fully counterbalanced. (b) Choice accuracy and RT are presented as functions of target values, distractor values, and distractor type (real vs. notional; within-participants factor). v1 < v2 < v3 < v4 indicating the four increasing value ranges. From trial to trial, the reward outcome was jittered with a uniform distribution. (c) Generalized Linear Model (GLM) is presented with the following variables: HV: higher-value target; LV: lower-value target; DV: distractor value; Dtype: a dummy variable with 0 for the notional distractor condition and 1 for the real distractor condition. Error bars represent ± s.e.m. across participants (N = 20). Asterisks denote significant effects (p < .05; two-sided one-sample t-tests against 0), following Holm’s sequential Bonferroni correction for multiple comparisons.

Upon fitting the learning model to these data, we found that its predictions were closely aligned with the observed behaviors (Fig. 4b). Inclusion of an additional divisive normalization component, however, resulted in incorrect behavioral patterns—particularly in the real distractor block, where the denominator is significantly inflated by a higher-value distractor (Fig. 4b: ‘Model’). Interestingly, we did observe a slowing in RT when confronted with a higher-value distractor, and this phenomenon was confined to the real distractor condition (Fig. 4b). This finding is consistent with previous research, which has shown an effect on RT but not on choice accuracy when a value-based distractor is present^24^. Taken together, these results further weaken the case for divisive normalization as a sole explanatory mechanism, while buttressing the role of learning asymmetry in shaping decision-making behavior.

### Magnitude sensitivity is nullified in the presence of counterfactual feedback

Lastly, we sought to substantiate the role of sampling-based asymmetry during learning as the source of observed decision bias. To this end, we explicitly neutralized key factors contributing to this asymmetry by providing participants with complete (full) feedback—revealing not just the outcome of their chosen option, but also of their unchosen one. This approach aimed to further elucidate the roots of the magnitude effect seen in Experiments 1 and 2, a phenomenon where choice accuracy improves as the overall value of the target options increases (Fig. 4b, c). For this purpose, we conducted a pre-registered Experiment 3 with 46 participants, employing a within-participants design that allowed for the comparison of learning and choice behavior under partial versus full feedback conditions across alternating blocks. Initial analyses revealed a replication of the positive magnitude effect—namely, that choice accuracy improved as the magnitude of targets rose—in the partial-feedback condition (Fig. 5a). Strikingly, the type of feedback wielded significant influence over this magnitude effect, i.e., the interaction term (HV + LV) x C, *t*(45) = -2.51, *p* = .016 (Fig. 5b; left). We found that the influence of target magnitude on choice accuracy effectively vanished under full-feedback conditions, *t*(45) = .87, *p* > .1. Moreover, reaction times (RT) displayed a declining trend as the value magnitude of the targets escalated (Fig. 5a, b): main effect of HV + LV on RT, *t*(45) = -4.93, *p* < .0001, and not influenced by feedback type (*p* > .1). Taken together, these findings bolster the notion that the provision of counterfactual feedback effectively nullifies the need for exploratory choices. This, in turn, ameliorates the impact of sampling-based learning asymmetry, such as the over-exploitation of high-value options that hinders effective learning of other available options. Thus, these data provide compelling evidence that sampling-based asymmetries during learning are a central mechanism driving observed decision biases, as opposed to divisive normalization or other competing explanations.

**Figure 5.**
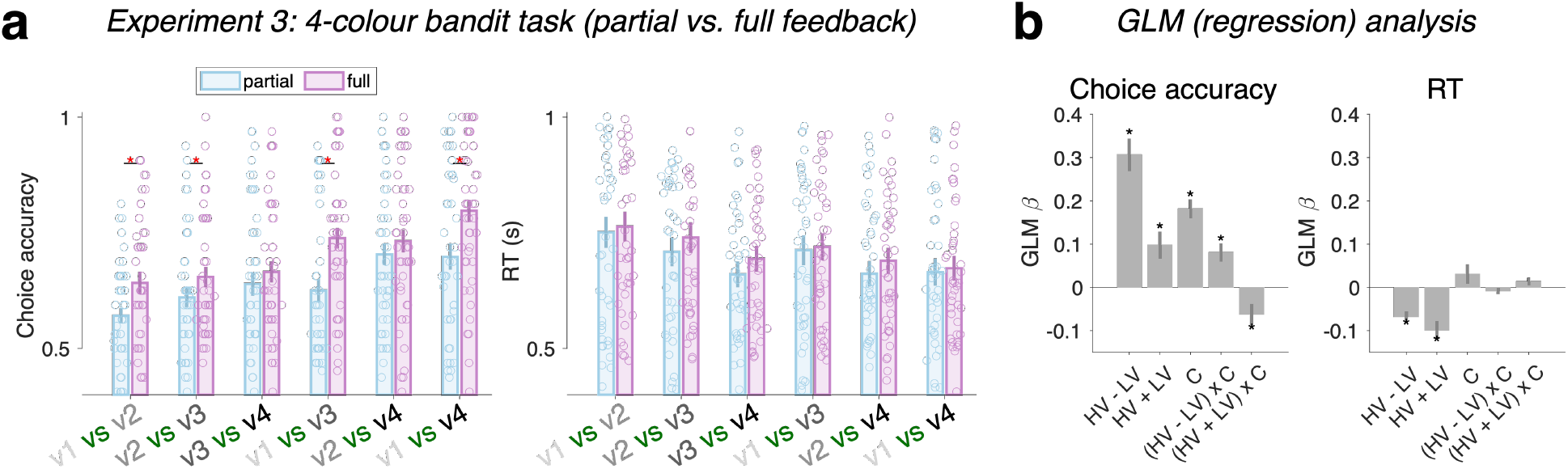
Behavioral data from Experiment 3 with counterfactual feedback. (a) In a pre-registered online Experiment 3, participants learned to choose between a left and a right stimulus, represented as abstract colorful fractals, without the presence of a distractor. The reward scheme paralleled that of Experiment 2, with four different fractals mapped onto four increasing value ranges (v1 < v2 < v3 < v4). ‘v1 vs v2’ denotes the condition where one target is associated with a lower value (v2) and the other with a higher value (v2), and the accuracy indicates the probability of choosing v2. ‘Partial’ refers to showing the reward outcome only for the chosen option, while ‘Full’ signifies showing the reward outcome for both the chosen and foregone options (i.e., counterfactual). (b) A Generalized Linear Model (GLM) is presented, incorporating the following variables: HV: higher-value target; LV: lower-value target; C (condition): a dummy variable with 0 assigned to the partial-feedback condition and 1 to the counterfactual-feedback condition. Error bars indicate ± s.e.m. across participants (*N* = 46). Asterisks denote significant effects (p < .05 two-sided one-sample t-tests against 0), following Holm’s sequential Bonferroni correction for multiple comparisons.

## Discussion

Contextual effects question the tenets of rational choice theory and provide insights into the computational underpinnings of decision-making. Recent research on these effects diverges in their conclusions, with some studies showing a negative distractor effect^9^ while others, showing positive^19^ and interacting distractor effects^18^. In the decision sciences, models are similarly divided: some disregard average value, concentrating solely on value differences^13^, while others posit a diminishing sensitivity to value^10,11^, and the negative effect of overall value magnitude of the context^32^. Challenging these diverse theories, new empirical data suggest that high-value options are more easily distinguishable, i.e., positive magnitude effect^34^. This contradicts predictions made by principles like diminishing value sensitivity and divisive normalization^9^, thereby offers a new piece of the puzzle by suggesting that distortions in value representation, previously attributed solely to hard-wired neural coding properties, are not the only factors contributing to contextual effects on economic decisions. Here, we studied the ways humans adjust their information-sampling strategies across diverse reward distributions during reward-guided learning. We obtained converging evidence for the presence of ostensible context effects, namely magnitude sensitivity and distractor effects. Crucially, these effects are not rooted in representational biases. Rather, they surface as incidental by-products of asymmetric sampling, which occurs in response to uneven reward distributions across alternatives.

The lack of value-based distractor effect we report here adds to extant replication failures of negative^19^ or interacting^18^ distractor effects in choice^24,26,28^. Notably, Gluth et al.^24,28^ claimed evidence for a distractor effect that influences only response times but not choice accuracy. Unlike the choice distractor effect, we here confirm that this effect on response time is not artefactual and not driven by the sampling asymmetry, as this effect is absent in the notional distractor context (Fig. 4b). Presumably this response-time effect reflects the fact that control processes are engaged more heavily to inhibit invalid motor responses towards a high-valued distractor. Whether this slow-down, represents a reduction in the value difference of the non-distractor alternatives alongside a compensatory adjustment of the speed-accuracy trade-off, or merely a change in ‘non-decision’ time remains an open question.

Storing information in an absolute form across multiple alternatives or attributes is not only inefficient^12^ but also falls prey to the challenges posed by the curse of dimensionality^8,40^. One plausible way to mitigate these issues is by selectively discarding less crucial information^41^. While this strategy aims for expediency, it frequently results in the over-exploitation of high-value options. In line with this spirit, strategy such as Selective Integration^27,42^ prioritizes larger values at the expense of smaller ones, resulting in a sampling asymmetry during the reward learning process. Such a selective approach may enhance robustness against decision noise^27,43,44^—a facet not explored in the current paper but warranting future investigation. From a normative theoretical perspective, the observed asymmetric sampling bias could be interpreted as a manifestation of a ‘winner-takes-all’ greediness^43^, arising from limitations imposed by finite cognitive resources^45,46^. This bias has been substantiated across a variety of task scenarios, serving as a normative rationale to counter computational noise^46^—the stochastic fluctuations affecting the internal world model^47,48^. Seminal research has blurred the lines between variance and bias^47^, while other influential studies have highlighted the advantageous role computational noise plays in mitigating rigidities and inflexibilities in reward-guided learning. Our study has not yet delved into the potential counteracting effects of learning noise in balancing the adverse impacts of sampling asymmetry. The current task design, nevertheless, presents an opportunity to fit noisy learning models and test their predictions. For example, the absence of feedback in probabilistic rewards would result in larger jumps in trial-to-trial reward prediction errors for higher-value options compared to lower-value options. In the realm of noisy value learning, the noise corrupting value updates scales multiplicatively with prediction error^49^, thereby counteracting the over-exploitation of lower-value options and rendering their value updates more precise. Future research should focus on individual differences in learning noise and examine their relationship with the extent of over-sampling bias, particularly in counterfactual feedback conditions where learning noise has been found to dominate decision variability^49^.

In conclusion, our research adds a critical voice to the ongoing challenge^50–53^ against the standard tenet of neuroeconomics that valuation is menu-invariant^54–56^. We highlight the importance of incorporating strategic learning dynamics to reconceptualize how biological brains strive to optimize decisions^57^. This becomes particularly significant when value is not anchored to sensory properties dictated by hardwired neural coding principles but is rather a dynamic construct continuously updated by the influx of newly sampled information.

## Methods

### Task and design

Across three experiments, either conducted in a lab or online, we recruited a total of 100 human participants to engage in variations of a multi-armed bandit task. In this task, different visual stimuli were associated with varying magnitudes of reward points. Participants were instructed to select either the left or right stimulus to maximize their total reward. They were advised to consistently opt for the stimulus offering a relatively higher reward in each trial. Importantly, no information regarding the mapping of colors to rewards was provided at the beginning of each new block. Furthermore, each new block featured a unique color-reward mapping, requiring participants to adapt and learn through a process of trial and error. The study was approved by the ethical committee responsible for the University Medical Center Hamburg-Eppendorf.

In Experiment 1 (N = 20, age range 20 to 36), on each trial, three iso-eccentric stimuli were presented, each rendered in a different color (red, green, and blue). Two target stimuli to the left and right sides of a central fixation point were available for choice (button press) whilst a third ‘distractor’ stimulus below the fixation was unavailable to choose. Each color was associated with a reward magnitude (reward p = 0.5), selected (with repetition) from a set of increasing values (1, 5, and 9 cents), and the color-reward mapping was fixed within a block but was fully counterbalanced across blocks. Human participants were instructed to learn the color value by trial-and-error and maximise cumulative rewards over trials. Importantly, in alternating blocks, the distractor was either rendered in color or replaced by a white circle (‘notional distractor’). Each block consisted of 36 trials, designed to fully counterbalance the combinations of color and location. Overall, participants completed a total of 54 blocks; half of these were designated as real distractor blocks and the other half as notional distractor blocks. The sequence of these blocks was randomized for each participant. For both types of distractor blocks—real and notional—the study employed a comprehensive counterbalancing scheme for color-reward mapping. Given that there were three colors to be mapped to three numerical values (1, 5, 9), and allowing for repetition, the total number of unique combinations amounted to 27 for each type of distractor block.

In Experiment 2 (N = 20, age range 20 to 40), we mapped four different colors to four reward levels (1, 2, 3, and 4) without repetition. Within each block, the 4 reward levels were: 10-18, 37-45, 64-72, 91-99, i.e., the bandwidth was 8 (thus no overlapping), and we jittered the trial-to-trial reward feedback using a uniform noise. Each block consisted of 48 trials, designed to counterbalance color-location combinations. In total, participants completed 16 blocks, with half categorized as real distractor blocks and the remaining as notional distractor blocks. A full counterbalancing scheme for color-reward mapping across blocks was not implemented, as it would have excessively prolonged the experiment. This decision was justified by our primary focus on within-participant comparisons between real and notional distractor blocks. Instead, we employed a balanced Latin-square design to generate four mappings, which were derived from two randomly selected seed mappings. This approach was applied independently for each participant, ensuring a high level of randomization.

Experiment 3 was performed online. On each trial, two visual stimuli (colorful fractal images^58^: http://www.schapirolab.org/resources) were presented on the screen, one to the left and the other to the right side of a central fixation “+”. In the instruction and practice, the participants were told to think of the stimuli as Casino tokens, which were worth different reward points. Participants were instructed to choose the token with a relatively higher reward on every trial. The experiment was divided into 8 short blocks (2.5 mins each). Each block contained a fixed number of trials (n = 48). Each block had a unique set of 4 fractals as stimuli; therefore, each block required learning the option values from scratch. On each trial, two different fractals were shown as the choice targets. In total, among the 48 trials per block, we factorially manipulated the stimuli combination (4C2 = 6) x location (2) x reward scheme (4 possible trial versions). The reward scheme was designed in a way to achieve 50% reward probability. That is, for instance, for a condition {A1, A2} where the left stimulus was A1 (true hidden reward: R1) and the right stimulus was A2 (reward: R2), there were 4 repeated trials with 4 different possible reward contingency rules: {R1, R2}, {0, R2}, {R1, 0}, {0, 0}. This design ensured that even if a participant always chose the left stimulus A1 in this condition, the chance of receiving the true reward “R1” was only 50% across these 4 repeated trials, and the other 50% of the times the participant got 0 as the reward outcome. Among the 8 blocks, half (4 blocks) were ‘partial feedback’ and the other half ‘full feedback’. For a partial feedback block, only the outcome of the chosen stimulus was displayed, whereas for a full feedback block, the outcomes of both stimuli were displayed. The feedback was displayed below the visual stimuli. Participants had 3 seconds max. to respond; otherwise, a warning sign “too slow!” was shown below the fixation cue. If a choice was made within the 3 s limit as required, the feedback was shown for 1 second (while the visual stimuli remained on the screen), and then the screen was cleared to get ready for the next trial. Within each block, the 4 reward levels were: 10-18, 37-45, 64-72, 91-99, i.e., the bandwidth was 8 (thus no overlapping), and we jittered the trial-to-trial reward feedback using a uniform noise. The jitter only applied to the true reward outcome, not when reward = 0 due to the above-mentioned probabilistic reward scheme. At the end of each finished block, the participants were shown the total reward points they had obtained in this past block out of xxx max. possible. To enable the analysis of noise decomposition used in a previous study^49^, we made 2 full-feedback blocks as the exact ‘twins’ of the other 2 full-feedback blocks (also called ‘replay’ blocks). This design could only be applied to the full-feedback blocks because this analysis required that the human choice variability didn’t impact the expected value updates in the learning model. Specifically, if we denoted the 8 blocks as A (partial), B (full), C (partial), D (full), E (partial), F (full), G (partial), H (full), then we made a replica of the exact trial order in B and assigned this order to block F, and in the same spirit, did this for D and H. So, B and F were twins, and D and H were twins, in terms of the trial-by-trial reward scheme and everything else except for the visual stimuli.

We generated the trial orders pseudo-randomly offline (in service of the ‘twin’ or ‘replay’ design), and for every 5 participants on Prolific, we used a new trial order, to achieve enough randomness across participants (target sample size: N = 45). For every participant, the ‘block’ order was completely randomized online by Gorilla; that is, the partial vs. full feedback block order was random across any participants. The task was built with Gorilla Experiment Builder, and participants were recruited via Prolific. The study took about 30 ∼ 40 minutes to complete. If a participant finished the study, they received £5 for their time. In addition, if they finished the study and we approved the submission, they could win a bonus up to £2 according to their task performance. After the 8 blocks, one of the blocks was selected randomly by us, and the participant received the bonus associated with this block. The bonuses associated with the other blocks were not considered; it was thus in their best interest to win as many points as possible on every block to increase the chances of leaving with a bonus. We did this bonus calculation offline by ourselves. Our target sample size was 45 human participants, but we recruited 60 participants because of the attrition/dropout rates during online recruitment. We estimated our target sample size based on a power analysis with an effect size of Cohen’s d = 0.5, alpha error = 0.05, power = 0.95, for a one-sided one-sample (paired-sample also) t-test. This power analysis yielded a sample size of 45 participants (calculated with G*Power 3.1^59^). This one-sample t-test examined whether the linear slope regressing choice accuracy against the option value level was above zero; that is, whether the choice accuracy between value level 1 vs. 2 was lower than the accuracy between level 2 vs. 3 and lower than that between 3 vs. 4 (one-sided test since our hypothesis was directed). We launched 12 batches on Prolific (5 participants per batch). After collecting the data from 60 participants, we checked whether we had at least 45 participants with full datasets (8 blocks were completed) that met the basic requirements for data quality and inclusion (see below). If that was not the case, then data collection proceeded until 45 eligible participants with full datasets had been acquired. The final dataset contained 46 participants. We pre-registered this study before data collection (see full pre-registration report on OSF: https://osf.io/nq532).

### Computational modeling

In all our tasks, reinforcement learning (RL) theory guided our modeling of participants’ behavioral adaptation for reward maximization. Within the RL framework, agents construct a probabilistic policy that prescribes the action to be taken in each state^7^. This policy principally relies on action-value functions, which encapsulate the expected reward for each action in the state. Agents refine these action values through ongoing observations of the rewards they receive at each time point t. The ‘reward prediction error,’ calculated as the discrepancy between the estimated reward Qt and the actual reward rt, serves to update these values, i.e., a delta learning rule. The weight assigned to prior value estimates versus new outcomes is modulated by the learning rate parameter α. Lower values of α emphasize past experiences, fostering stable learning across extended time spans. Conversely, higher α values prioritize new outcomes, facilitating more rapid and adaptive behavior over shorter intervals. To translate action values into actionable policies, most cognitive RL models employ a ‘softmax’ function, which introduces a level of stochasticity into the decision-making process^4^.

Prior research indicates that humans exhibit differential learning rates for positive and negative prediction errors, suggesting the presence of asymmetric learning mechanisms^36–38^. In addition to utilizing the standard delta rule, which employs a single learning rate, we also explored the potential for dual learning rates that separately account for positive and negative reward prediction errors^39,60^. Beyond the basic updating of action values, or Q-values, these can also be normalized by the sum of the values corresponding to all presented options (Fig. 3). This additional normalization step offers to assess its alignment with observed patterns of human behavior. The Q value at the beginning of each block was set to a free parameter. Models were fit to data via maximising the log-likelihood summed over trials. Because the *LL* functions all have analytical solutions, Matlab’s fmincon was used for parameter optimisation. To avoid local optima, we refit each model to each participant’s behavioural data at least 10 times using a grid of randomly generated starting values for the free parameters. Conversative criteria were used during the optimisation search: MaxFunEvals = MaxIter = 5,000; TolFun = TolX = 10^−10^.

## Acknowledgements

This work was supported by the EU Horizon 2020 Research and Innovation Program (ERC starting grant no. 802905) to K.T. The funders had no role in study design, data collection and analysis, decision to publish or preparation of the manuscript. We extend our gratitude to M. Tohidi for assistance with data collection and to M. Siems for discussion during the early stage of this research.

